# Decoding intended speech with an intracortical brain-computer interface in a person with longstanding anarthria and locked-in syndrome

**DOI:** 10.1101/2025.08.12.668516

**Authors:** Justin J. Jude, Stephanie Haro, Hadar Levi-Aharoni, Hiroaki Hashimoto, Alexander J. Acosta, Nicholas S. Card, Maitreyee Wairagkar, David M. Brandman, Sergey D. Stavisky, Ziv M. Williams, Sydney S. Cash, John D. Simeral, Leigh R. Hochberg, Daniel B. Rubin

## Abstract

Intracortical brain-computer interfaces (iBCIs) for decoding intended speech have provided individuals with ALS and severe dysarthria an intuitive method for high-throughput communication. These advances have been demonstrated in individuals who are still able to vocalize and move speech articulators. Here, we decoded intended speech from an individual with longstanding anarthria, locked-in syndrome, and ventilator dependence due to advanced symptoms of ALS. We found that phonemes, words, and higher-order language units could be decoded well above chance. While sentence decoding accuracy was below that of demonstrations in participants with dysarthria, we are able to attain an extensive characterization of the neural signals underlying speech in a person with locked-in syndrome and through our results identify several directions for future improvement. These include closed-loop speech imagery training and decoding linguistic (rather than phonemic) units from neural signals in middle precentral gyrus. Overall, these results demonstrate that speech decoding from motor cortex may be feasible in people with anarthria and ventilator dependence. For individuals with longstanding anarthria, a purely phoneme-based decoding approach may lack the accuracy necessary to support independent use as a primary means of communication; however, additional linguistic information embedded within neural signals may provide a route to augment the performance of speech decoders.

---

Loss of communication is one of the most disabling and distressing symptoms of amyotrophic lateral sclerosis (ALS), brainstem stroke, and other neurologic conditions that cause paralysis. The ability to communicate with depth and nuance is one of the most defining characteristics of humans, and for people with ALS, the potential loss of communication is often a determining factor in the decision to continue or withdraw life-sustaining care ^1^. Commercially available augmentative and assistive devices may enable maintained communication by relying upon residual motor control (e.g., eye gaze) but they are often slow, error-prone, and difficult to use, leading to high rates of abandonment ^2,3^. For people with ALS, stroke, muscular dystrophy, and other neurological conditions who have lost or are losing dexterous control of their hands and their ability to speak, there is an urgent need to provide intuitive, accurate, and facile tools to restore and maintain communication.

Intracortical brain computer interfaces (iBCIs) ^4,5^ restore lost function by extracting information directly from the cerebral cortex, thereby bypassing the site of primary pathology. Previously, iBCIs have been used to restore communication for people with paralysis by providing point-and-click control of a digital interface ^6–10^, such as a computer cursor that can be used to select letters from a virtual on-screen keyboard. Though these systems have the advantage of rapid calibration ^11,12^, their overall communication rates are limited by the speed at which the user can click on individual letters on a computer screen.

More recently, iBCIs and electrocorticography (ECoG)-based BCI systems have been shown to be effective at higher throughput communication, for example through articulator-based speech decoding ^13–26^, character-based handwriting decoding ^27,28^, and finger-based typing ^29,30^. In these systems, neural activity pertaining to the intended movement, in the absence of actual movement of speech articulators or hand effectors, can be used to decode rapid sequences of phonemes or characters to construct language. High accuracy in recent work ^14–16,27,29^ is maintained through the use of language models ^31^ that take sequences of predicted phonemes or characters and select the most likely intended output sequence based on statistics of large English language datasets ^32^. Similarly to recent work ^16^, we displayed decoded words on a screen as they were decoded, after which a text-to-speech model ^33^ was used to vocalize each decoded sentence in a voice similar to the participant’s original voice before ALS onset.

Prior research has shown that speech iBCIs can be used by people with dysarthria and other conditions that cause articulator paralysis to communicate accurately and with high throughput ^14–16^. Recent studies in particular demonstrate rapid calibration of such a system, achieving highly accurate communication within a single day ^16^ and prolonged independent use of a speech neuroprosthesis for several months ^18^. In this work, we show that a speech iBCI neuroprosthesis can be used by a person with longstanding anarthria and locked-in syndrome. We report on the possibilities and challenges of developing a speech neuroprosthesis for an individual who has not spoken in over two years.

Previous work on speech decoding in anarthric participants has primarily involved ECoG recordings ^34^. This includes single word decoding constrained to a 50 word vocabulary ^35^ and more recently, large vocabulary sentence decoding ^14^. A previous attempt at communication with a locked-in person with advanced ALS, similar to our study participant, has demonstrated the feasibility of using an implanted BCI system for long-term, reliable communication, but throughput was limited to binary classification ^36,37^. In contrast, here we assess speech decoding performance with a comprehensive and thus much more pragmatically useful unconstrained vocabulary.

In this work, we explored the feasibility of speech decoding in iBCI research participant T17, who has locked-in syndrome. As the participant is anarthric, his enrollment in the clinical trial permitted an opportunity to assess whether articulator control is sufficiently represented in the underlying neural activity in ventral, middle and dorsal precentral gyrus of a person that has been unable to speak for more than two years. We posit that, although neural activity pertaining to orofacial movements persisted, control of behavior specifically pertaining to fine articulator movement for phoneme production may not have been completely preserved.

In this case, articulator-based decoding from ventral precentral gyrus caused several instances of overlapping neural signatures during generation of phonemes that entail combinations of similar orofacial movements. Given this barrier for participants with longstanding anarthria, we additionally present evidence demonstrating the feasibility of integrating the decoding of larger language units such as phrases and full sentences from middle precentral gyrus into a real-time phonemic speech decoding pipeline to augment performance.

## Results

### Neural tuning to phonemes varies along precentral gyrus

As a first assessment of neural population tuning, we asked participant T17 (Fig. 1a) to perform a series of attempted isolated facial movements and phoneme articulations. In an instructed delay paradigm, individual trials of 13 distinct orofacial movements and 39 unique phonemes were cued (Fig. 1b). Both arrays placed in area 6v had a similar number of strongly tuned (> 0.5 fraction of variance accounted for (FVAF)) electrodes to the 13 orofacial movements, while the more dorsal of the two 6v arrays had more strongly tuned electrodes to the 39 phoneme articulations (Fig 1c). This is contrary to the tuning in 6v arrays found in another clinical trial participant (T12) with intracortical arrays ^15^, where the more dorsal 6v array electrodes were more tuned to orofacial movement conditions than phonemes and vice versa for the ventral 6v array.

**Figure 1:**
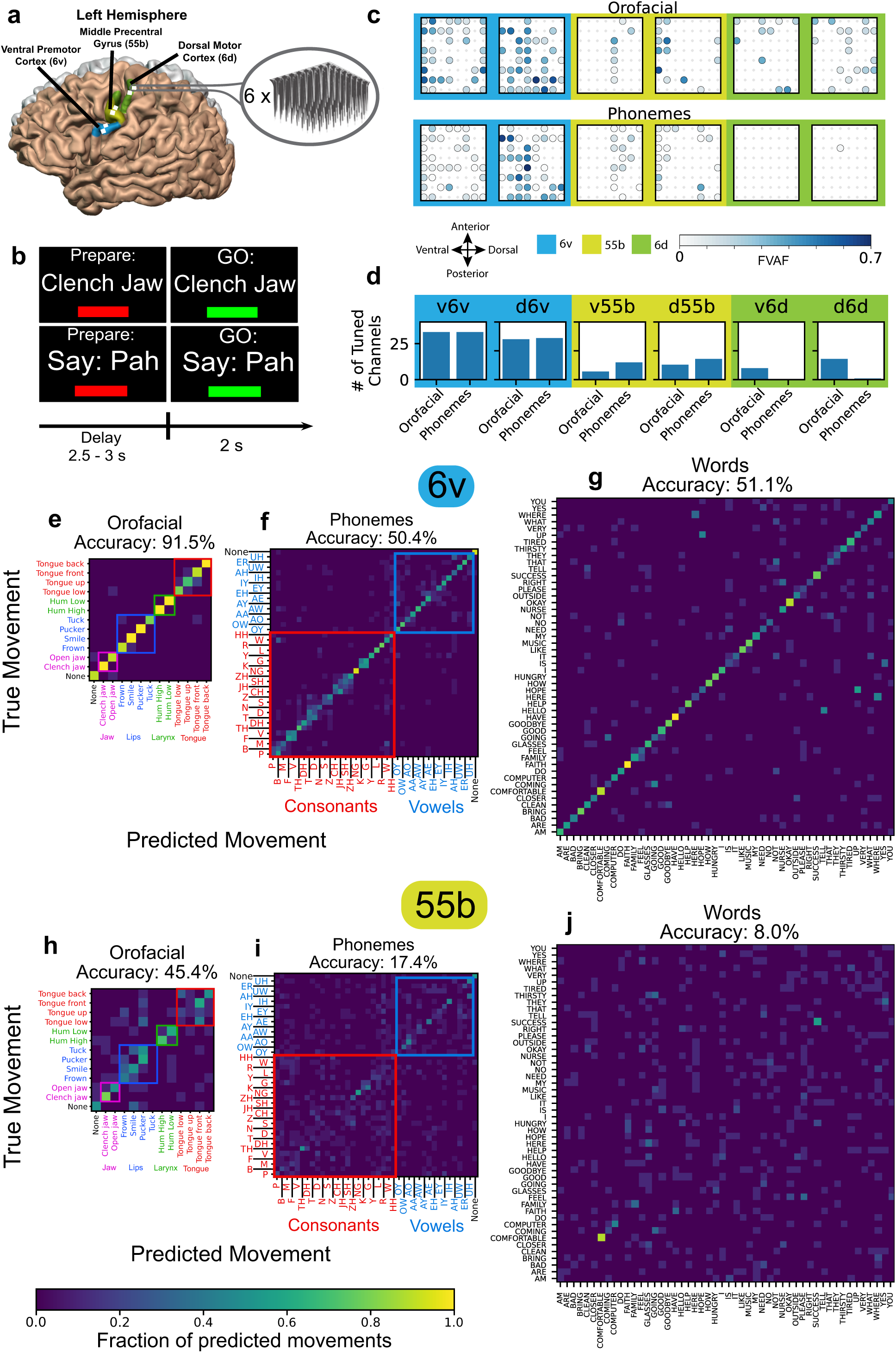
Area 6v encodes orofacial movements and phonemes in a locked-in participant. **a**, Participant T17 microelectrode array locations in the left hemisphere. Two arrays were placed in each of the following cortical areas: dorsal (6d), middle (55b) and ventral (6v) precentral gyrus. **b**, Instructed delay isolated cue task: the participant was instructed to attempt to speak a cued word (either with a single phoneme or a word with multiple phonemes) (top) or perform an orofacial movement (bottom). **c**, Tuning heatmaps for all six arrays; backgrounds are shaded by cortical area. Circles indicate that threshold crossing features on a given electrode varied significantly across all orofacial movements (top) or all phonemes (bottom) (p < 1 × 10^−5^ assessed with one-way analysis of variance). Shading indicates the fraction of variance accounted for (FVAF) by cross movement or cross phoneme differences in threshold crossing rate. **d**, Number of tuned electrodes where threshold crossing features on a given electrode varied significantly across all orofacial movements (top) and all spoken phonemes (bottom) (p < 1 × 10^−5^ assessed with one-way analysis of variance). **e,f,g**, Confusion matrices showing decoding errors when using a simple Gaussian Naive Bayes classifier to predict **e**, isolated orofacial movements, **f**, isolated phonemes, or **g**, whole individual words from threshold crossing neural features from the 128 intracortical electrodes implanted in area 6v. **h,i,j**, Similarly, decoding accuracy using threshold crossing features from the 128 intracortical electrodes implanted in area 55b.

The area 55b arrays had 3 electrodes strongly tuned to orofacial movements (> 0.5 FVAF), especially in the dorsal 55b array, while neither area 55b array had strongly tuned electrodes to the 39 phoneme articulations. Far fewer of T17’s 55b electrodes were tuned to either orofacial movements or phonemes compared to electrodes in T17’s 6v arrays, although the average threshold crossing rate was lower overall in electrodes in the 55b arrays (Supplementary Fig. 1). A small number of electrodes in area 6d were weakly (< 0.2 FVAF) to moderately (> 0.2 and < 0.5 FVAF) tuned to the orofacial movements, with one electrode being weakly tuned to phoneme articulation ^38,39^. We observed the largest number of overall tuned single electrodes to conditions of both tasks in the ventral 6v array (Fig. 1d).

The high number of tuned electrodes in area 6v yielded high decoding accuracy when using a simple linear Gaussian Naive Bayes (GNB) classifier (with leave-one-out evaluation) to predict orofacial movements (91.5% (95% CI=[88.4, 94.6])) (Fig. 1e), phonemes (50.4% (95% CI=[48.1, 52.8])) (Fig. 1f) using just these 128 electrodes. Notably, decoding accuracy of orofacial movements is significantly higher than decoding accuracy of phonemes (p < 5.57 × 10^−19^ assessed with one-way analysis of variance) compared to respective chance levels. Moreover, similarly articulated consonant phonemes are less distinguishable by the decoder than with vowel phonemes, for example between labial phonemes such as P and B and between alveolar phonemes such as CH and JH. Similarly, individual vowels were most often confused with other vowels rather than consonants.

When decoding single words from a 50 word vocabulary ^13^, we found well above chance accuracy (51.1% (95% CI=[48.0, 54.2])) using a GNB discriminator and threshold crossing features from area 6v electrodes (Fig. 1g). This higher accuracy compared to isolated phoneme decoding (relative to chance) may be due to neural activity pertaining to single words, i.e., to a combination of several phonemes in a limited vocabulary, being somewhat easily separable, especially when these phoneme combinations are unique. Most word classification errors when decoding from area 6v electrodes arise from words with overlapping phonemes, such as “where” and “here” as well as “hello” and “tell”, which may suggest that the unit encoded in 6v is primarily phonemic or motoric as opposed to encoding entire words or larger language units such as phrases or sentences.

In area 55b, there were relatively fewer and less strongly tuned electrodes to orofacial movements and phonemes, resulting in lower decoding accuracy in both instances (Fig. 1i), 45.4% (95% CI=[40.0, 50.8]) and 17.4% (95% CI=[15.6, 19.2]) respectively. However, decoding of some orofacial movements, such as jaw and larynx-based movements (Fig. 1h), was reliably accurate. Additionally, we noted that longer length words such as “comfortable”, “computer” and “success” have the highest decoding accuracy using 55b area electrodes (Fig. 1j), suggesting a potential role in encoding larger language units, as we explore below.

### 6v and 55b neural activity cluster according to articulatory movements and to components of speech respectively

To further explore the distribution of neural activity underlying the population tuning in these cortical areas, we calculated the mean Mahalanobis distance ^40,41^ (a distance metric in multivariate space) between neural activity recorded during each attempted phoneme or orofacial gesture. We used these distances to run an unsupervised clustering algorithm yielding hierarchical dendrograms of neural tuning. We found that for area 6v, neural activity from the isolated phoneme task was neatly clustered by tongue position within the articulatory tract during phoneme production (Fig. 2a). Groups of bilabial, labiodental, alveolar, dental, rhotic, and post-alveolar phonemes were each clustered together, with vowels and consonants distributed throughout the dendrogram.

**Figure 2:**
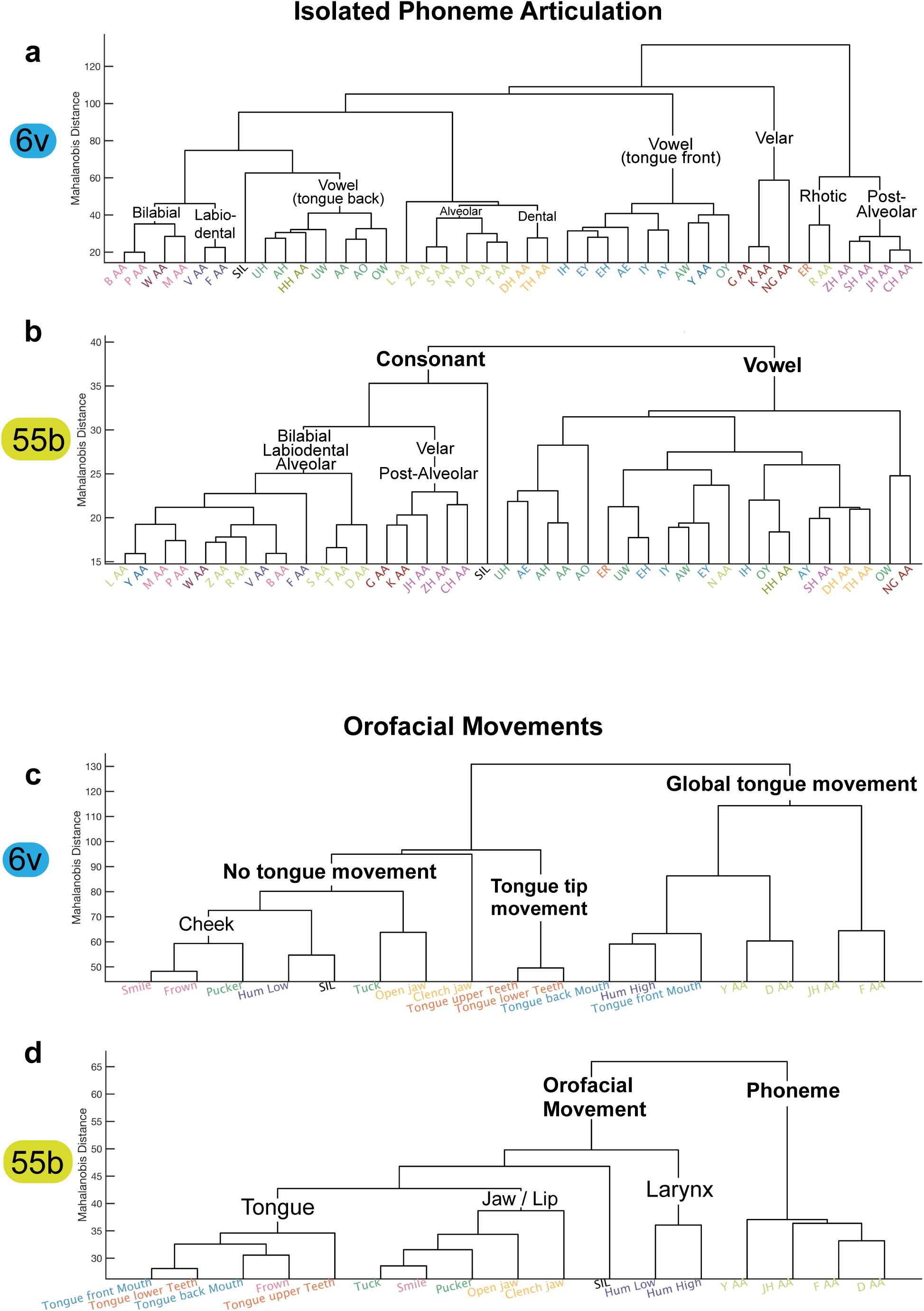
Phonemic and orofacial representations cluster differently depending on array location in PCG. **a**, Dendrogram based on the distance of neural activity recorded from the two 64 array electrode arrays in area 6v during the isolated phoneme production task. For area 6v activity, clustering depends most strongly on position of the tongue (i.e., phoneme place) within the articulatory tract during attempted phoneme production. **b**, For area 55b electrode neural activity, phonemes are most prominently clustered into consonants versus vowels, suggesting higher order classification. **c**, For the isolated orofacial movement task (which included twelve orofacial gestures and four phonemes), area 6v again demonstrates clustering based on tongue position. **d**, In area 55b, phonemes are clustered separately from the other orofacial movements, again suggesting stronger encoding of language than specific articulatory movements.

In contrast, the clustering of neural activity from area 55b most prominently separated consonants from vowels but otherwise was relatively insensitive to phoneme place (location of sound production in the mouth) or manner (how a sound is produced) (Fig. 2b). Similarly, when examining the neural activity from the orofacial movement task, activity from area 6v was again clustered most prominently by tongue position (Fig. 2c). Interestingly, in area 55b, neural activity from the orofacial gesture task (which included four phoneme cues along with twelve other non-language cues) was clustered most prominently between phonemes and non-language orofacial gesture cues (Fig. 2d).

### Sentence speech decoding

We next asked participant T17 to attempt to speak whole sentences cued on screen (Fig. 3a). Threshold crossing and spike power features from both area 6v and both area 55b microelectrode arrays (a total of 256 electrodes) were used to decode phonemes every 80ms using a recurrent neural network (RNN). Similarly to recent work ^15,16^, we used a 5-gram language model performing Viterbi (beam) search to infer the most likely sentence spoken given the complete sequence of RNN phoneme probabilities at all timesteps and the statistics of the English language (see Methods for further decoding pipeline details). Upon completion, each completed decoded sentence was synthesized into audio using a text-to-speech model ^33^ which was personalized to the participant.

**Figure 3:**
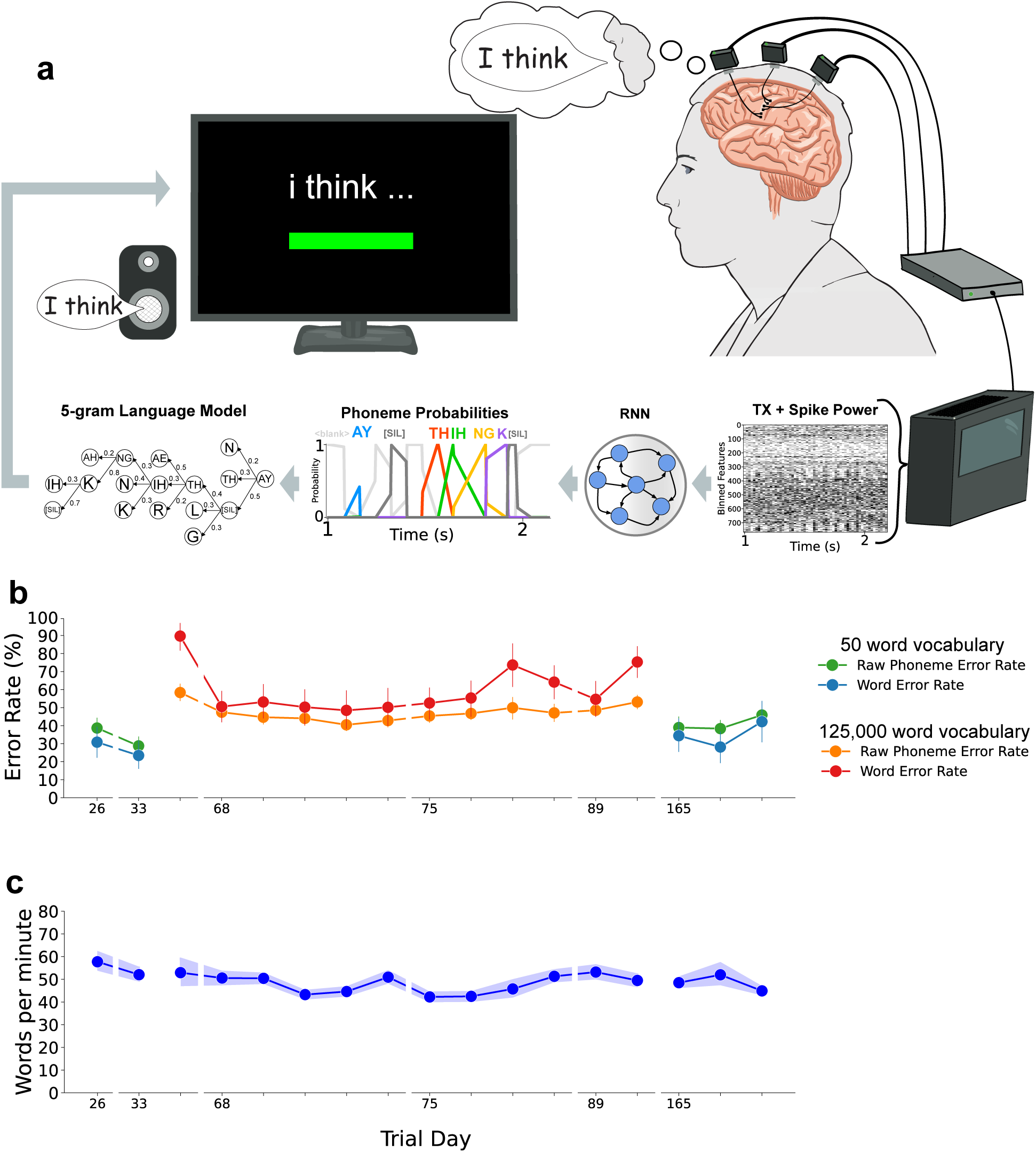
**a**, Sentence decoding pipeline - phoneme probabilities are inferred by a recurrent neural network every 80ms. Probabilities across all preceding time windows are considered by a 5-gram phoneme-based language model. The currently predicted sentence is displayed on screen. Each completed sentence decoded is synthesized into audio using a personalized text-to-speech model. **b**, Real-time decoded phoneme and word error rates across trial days, separated by trial days where sentence vocabulary is limited to 50 words (green and blue markers) and sentences consisting of an unconstrained vocabulary (red and yellow markers). Each tick represents a block on a given trial day. Error bars indicate a 95% confidence interval. **c**, Real-time speaking rate across trial days, measured in words per minute. Each tick represents a block on a given trial day. Shaded region indicates a 95% confidence interval.

In a sentence copy task, participant T17 was instructed to speak cued sentences from a 50-word vocabulary sentence corpus created using words from recent work ^35^. These are relatively simple sentences designed to be useful for communication in a care facility setting. In the first session in which sentence speech was decoded using the above pipeline, raw phoneme error rate averaged 38%, falling to 27% in the second session. Accordingly, word error rate fell from an average of 30% in the first session to 22% in the second session (Fig. 3b).

Subsequently on trial day 33, we switched to sentence cues sourced from a conversational English corpus ^42^ with an unconstrained vocabulary size. We initially observed a high raw phoneme error rate (56%) and a subsequent high word error rate (89%) when decoding conversational English sentences online. On trial day 68, online decoding on the first block yielded a lower average online raw phoneme error rate of 48% with a 50% average word error rate. Phoneme and word error rate reached their lowest on the fourth real-time decoding block of trial day 68, with a raw average phoneme error rate of 41% and an average word error rate of 48%. Phoneme error rate thereafter stayed consistently below 50% for the conversational English sentences, with word error rate usually slightly higher than this, sometimes much higher as on trial day 75.

Speaking rate averaged between 45 and 60 words per minute across all sessions (Fig. 3c). Owing to the participant’s lack of volitional control of his articulators, all vocalized speech was attempted without any actual orofacial movement. Thus, the rate of speech was not limited by articulator movement as with prior studies ^14–16^. Nonetheless, the participant was asked to attempt to speak slower than a typical conversation speed of 150 words per minute to enable the decoder outlined in Fig. 3a to more reliably delineate between phonemes.

### Sources of error in phoneme decoding

Similar to previously reported studies ^15,16^, training decoders with an increasing number of spoken sentence trials collected over successive sessions decreased the average offline phoneme error rate (Fig. 4a). We found that training on closed-loop blocks, i.e., those where sentences are decoded and results presented in real-time, had an outsized effect on improving the phoneme error rate compared to training just on open-loop blocks recorded on the same day, bringing the overall phoneme (word) error rate down to 34% (39%) offline. This suggests that sentence trials from closed-loop blocks are more informative due to the feedback received by the participant, perhaps due to higher engagement with the interactive task.

**Figure 4:**
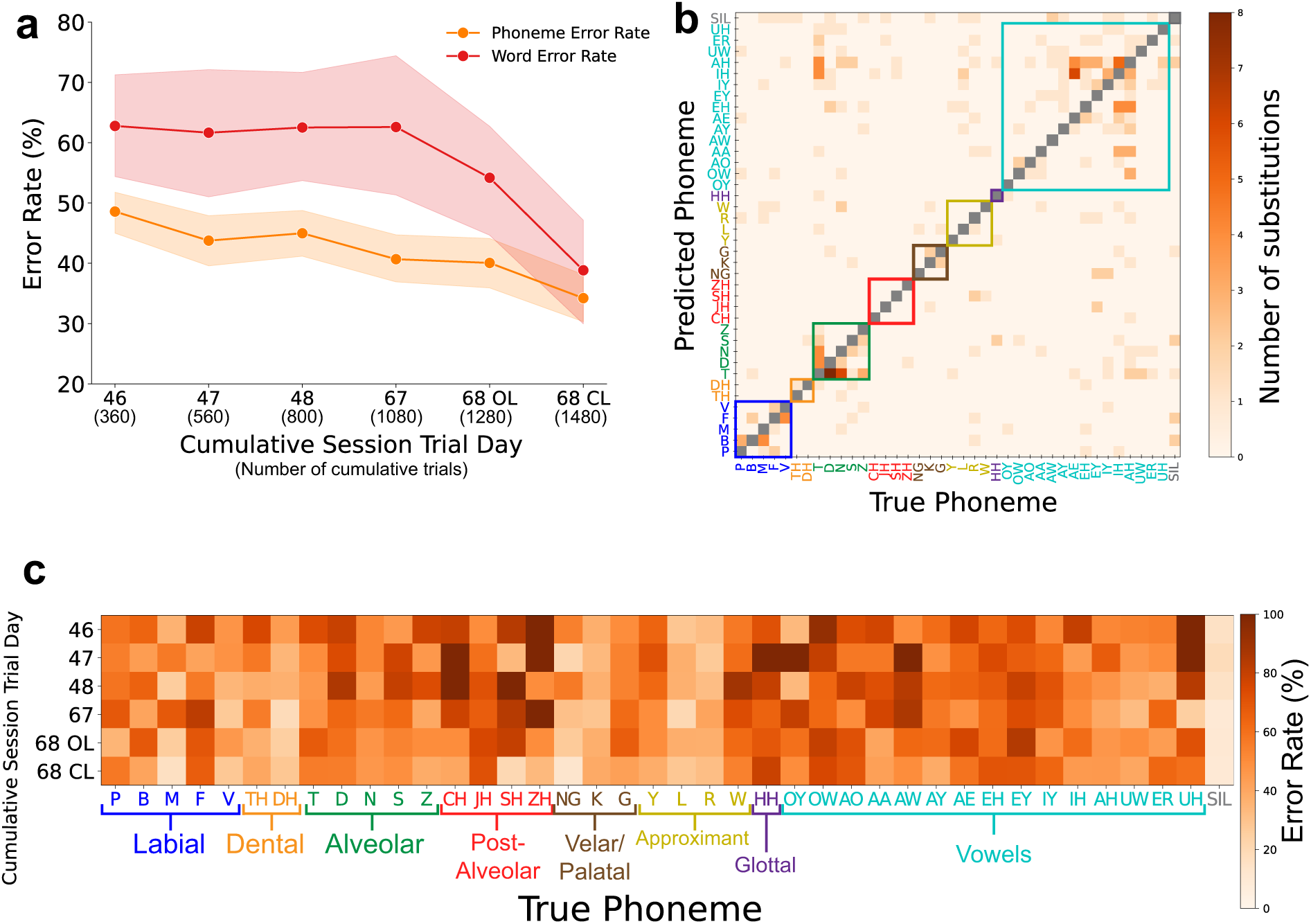
Training on sentences from several days improved decoding accuracy. Training on closed-loop data resulted in out-sized decoding improvement. **a**, Sentences from cumulative sessions were used to train a recurrent neural network (RNN) offline, with raw phoneme error rate and word error rate post language model reported with all data up to and including the listed session. OL indicates all open-loop sentences (no real-time decoding) and CL indicates the further inclusion of closed-loop sentences, where online feedback of decoded sentences was presented to the participant. Shaded regions indicate a 95% confidence interval. **b**, Substitutions required to produce ground truth phoneme sequences from inferred phoneme sequences, using an RNN decoder trained offline with sentences from 5 sessions. **c**, Heatmap showing individual offline average phoneme error rates across days during inference with an RNN trained up to and including sentence trials of each row’s trial day.

Interestingly, not all phonemes showed a similar improvement after RNN training with more data. Decoding of several consonants, such as M, DH, and NG, improved drastically when training on an increasing number of trials (Fig. 4c). Consonant phonemes such as SH and CH also had marked improvements in offline decoding accuracy, especially when the RNN was trained on closed-loop sentences. However, vowel phonemes did not improve substantially in accuracy with an increase in training sentences. Decoding of isolated phonemes (Fig. 1f) and decoding of phonemes from spoken sentences appear to have opposing confusion profiles, wherein, decoding of isolated phonemes caused greater confusion with consonants in isolated trials, whereas there was greater confusion with vowels in sentence decoding. This may be due to co-articulations during vowel production being more difficult to separate from adjacent phonemes when spoken in sequence, especially noting the participant’s longstanding anarthria.

To better understand the pattern of the decoding errors, we next examined offline substitution errors with an RNN decoder trained on trials from 5 sessions and tested on 50 held-out trials from the lattermost session. For consonant phonemes, sequence substitution errors were usually across similarly produced phonemes (e.g., V often substituted with F and D often substituted with T) (Fig. 4b). Similarly, with vowel phonemes most substitution errors were with other vowels. Particular vowels such as AH and IH were consistently predicted as other vowels and conversely other vowels were consistently predicted as AH and IH. However, there were several errors where consonants were predicted as vowels and vice versa, notably with alveolar consonants such as T and D.

### 6v and 55b encode language at different timescales

To better understand the functional localization in speech and language processing in the cortical areas from which we recorded, we conducted a three-phased reading, internal speech, and attempted vocalized speech task (Fig. 5a), similarly to previous work ^19,43^. On each trial, a phoneme, word, phrase or sentence (a “language unit”) was first cued to be read during the “reading phase”. After a two second delay, the participant was then cued to say the language unit to himself from memory without vocalizing in the “internal speech” phase; and after another two second delay, he was instructed to then attempt to vocalize the language unit from memory in the “attempted speech” phase. Trials consisted of 5 repetitions of 10 unique conditions of each language unit (phoneme, word, phrase, sentence), for a total of 40 unique stimuli (see Supplementary Table 3 for cue list). Stimuli from each language unit were of a similar length. Two of the ten word and two of the ten phrase stimuli were nonsensical (e.g., the nonsense word stimuli were “zelwog” and “cheldgup”).

**Figure 5:**
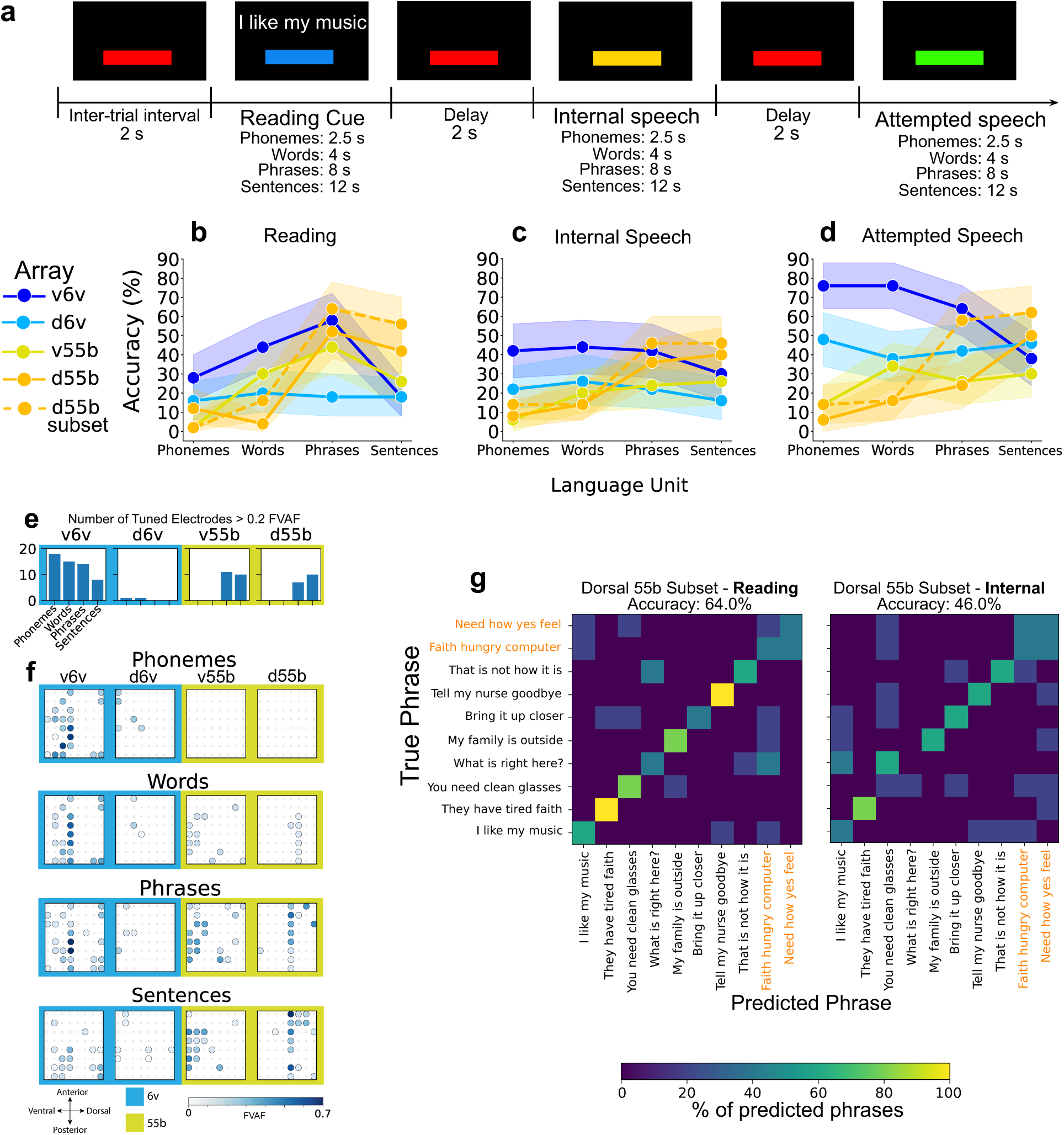
55b encodes phrase and sentence level units of language. **a**, In the three-phased task, each trial consisted of a reading, internal speech and attempted speech phase. **b,c,d**, Average decoding accuracy of phonemes, words, phrases and sentences in each of the **b**, reading, **c**, internal speech and **d**, attempted speech phases, colored by implanted array. Shaded regions indicate a 95% confidence interval. **e**, Number of tuned electrodes across all task phases for each language unit in each of the four arrays in speech areas. **f**, Tuning heatmaps for the four arrays in speech areas, with backgrounds shaded by cortical location (6v in blue, 55b in yellow). Circles indicate that threshold crossing features on a given electrode varied significantly across all task phases for a given language unit (phonemes, words, phrases or sentences) (p < 1 × 10^−5^ assessed with one-way analysis of variance). Shading indicates the fraction of variance accounted for (FVAF) by cross phase differences in threshold crossing rate. **g**, Confusion matrices showing decoding errors when decoding phrases in the reading (left) and imagined (right) phases of the task using a Gaussian Naive Bayes decoder on a subset ensemble of 20 contiguous electrodes from the dorsal 55b array. Nonsense phrase stimuli are highlighted in orange.

Across each phase of the task, neural decoding from the dorsal 6v electrode was relatively consistent across all four language units. Across all language units, average decoding accuracy increased from the reading phase (Fig. 5b) at 16%, to the internal speech phase (Fig. 5c) at 18%, to an average accuracy of 44% in the attempted speech phase (Fig. 5d) when decoded using a Gaussian Naive Bayes decoder. In contrast, we found that the ventral 6v array electrodes contained information preferentially about shorter units of language: neural activity measured from the ventral 6v array could decode phonemes, words, and phrases well (phrases with greater than 64% accuracy during the attempted speech phase (Fig. 5d)), whereas sentence decoding was far less accurate. This relative pattern of lower sentence accuracy was observed across all three task phases.

Meanwhile, the area 55b arrays, and the dorsal 55b array in particular, appeared to encode the longer units of language, phrases and sentences (i.e., those with contextual information), much better than phonemes and words, especially during the reading phase (Fig. 5b). Notably, phoneme, word, and sentence average decoding accuracy from dorsal 55b was similar across all three phases, whereas average phrase decoding accuracy was highest in the reading phase at 52%, decreasing to 38% in the internal speech phase, and lastly decreasing to 24% in the attempted speech phase. Phrase decoding was higher in all phases with a contiguous subset of dorsal 55b array electrodes which had sustained activity across all task phases (including the delay periods) (Supplementary Fig. 2). The ventral 55b electrodes had similarly increased average accuracy when decoding words and phrases as opposed to phonemes in both the reading and internal speech phases, but, sentence decoding accuracy was lower, similar to that of the 6v arrays. Notably, across all three phases, average word decoding accuracy was higher with the ventral 55b electrodes than with the dorsal 55b electrodes.

Classification accuracy across language units was consistent with the number and strength of tuned electrodes within each array (Fig. 5e,f), regardless of task phase. Across all task phases, the ventral 6v array had strongly tuned (> 0.5 FVAF) electrodes encoding phonemes, words and phrases, with these electrodes only weakly tuned (< 0.2 FVAF) to sentences. This array had several moderately tuned electrodes encoding phonemes and words but these only weakly encode phrases. A few dorsal 6v array electrodes only weakly encode phonemes, with minimal tuning to words, phrases and sentences in the few electrodes encoding these language units. This is in contrast to the encoding of phonemes in the phoneme cued task (Fig. 1c) where the participant was asked only to attempt to vocalize across all 39 English phonemes. In that task, there are a high number of moderate to strongly tuned electrodes in the dorsal 6v array. In this task, tuning in individual electrodes in the dorsal 6v array was less apparent in the reading and internal speech phases (Supplementary Fig. 3a,b), consistent with lower decoding accuracy in these phases when decoding from the dorsal 6v array electrodes (Fig. 5b,c).

Conversely, the ventral and dorsal 55b arrays had no tuned electrodes to this subset of 10 English phonemes in any of the task phases (Supplementary Fig. 3a,b,c), with an increasing number and intensity of tuned electrodes as the language unit size increased (Fig. 5e,f). There was a noticeable increase in the number and intensity of tuned electrodes encoding phrases in both 55b arrays across all task phases, consistent with the subset of contiguous electrodes in dorsal 55b with sustained activity across all task phases. There was a further strengthening of tuning to sentences in two of the electrodes tuned to phrases in the dorsal 55b array while almost all electrodes tuned to phrases in the ventral 55b array were also similarly tuned to sentences.

Looking more closely at decoding errors when decoding using a GNB decoder with the contiguous subset of electrodes with sustained activity in the dorsal 55b array (Supplementary Fig. 2), we note symmetrical confusion between the two nonsense phrases, “Need how yes feel” and “Faith hungry computer”, compared to sporadic errors for all other phrase pairs (Fig. 5g), in both the reading and imagined speech phase of the task. However, there was no notable confusion between the two nonsense word stimuli: “zelwog” and “cheldgup” (Supplementary Fig. 4). This suggests that these electrodes in the dorsal 55b array may encode sensical semantic context ^44^ at the phrase level rather than articulatory features.

## Discussion

The speech and motor impairments caused by ALS adversely affects the ability to effectively communicate. Recently, high performance speech iBCIs ^14–16^ have been demonstrated in dysarthric individuals with persistent phonation, however, the feasibility of such a system for individuals with complete and prolonged loss of motor control of the articulatory motor system has not previously been explored. Moreover, articulatory speech decoding had yet to be demonstrated in an individual with locked-in syndrome ^45^.

We found that decoding accuracy of isolated orofacial movements was significantly higher than that of isolated articulated phonemes compared to respective chance levels. This may be due to the longstanding anarthria of participant T17, causing a decline in the learned behavior corresponding to the production of the complex series of orofacial movements required to generate a differentiable neural signature for similarly articulated phonemes. Under this hypothesis, isolated movements were decoded with higher accuracy because these simpler movements were easier for the participant to attempt.

In this work we show speech sentence decoding with well above chance phoneme and word decoding accuracy in an iBCI clinical trial participant with longstanding anarthria and locked-in syndrome. Sentence-level speech decoding with a locked-in participant represents an advance compared to a fully implanted binary communication system using electrocorticography (ECoG) ^36,37^, albeit in only a copy-task and not implemented as a long-term independent means of communication. Importantly, real-time sentence decoding word error rate is above the word error rate shown with other participants ^14,15^ and above the word error rate that would be required for independent use for primary communication ^16,18^. In these recent studies ^14–16^, participants have been dysarthric but have maintained some volitional control of speech articulators, facial muscles, and muscles and apparatus of phonation. The improvement shown in this work in phoneme decoding accuracy when a decoder was trained on closed-loop sentences suggests a path toward improved articulator-based decoding with continued real-time feedback, though the extent to which such a learning-based strategy will help remains to be seen.

Electrodes in area 55b contributed greater word level accuracy to longer words, indicating higher than phoneme level language encoding. This is consistent with recent work outlining the role of area 55b in higher-order speech planning ^46–48^, suggesting that encoding of facial articulator and vocal tract muscle movements may be less prevalent than with area 6v. The phrase and sentence level encoding observed, which was especially prevalent in 55b, suggests that while cortical area 6v should be used to decode phonemes ^14–16^, neural activity from cortical area 55b could also be incorporated into decoding at a higher language level.

Future work will focus on a path to independent use of such a system, even when phoneme decoding from intracortical electrodes implanted in speech motor areas such as 6v is suboptimal. Although real-time feedback may improve articulator based decoding, we hypothesize that higher level encoding of intended language output at the phrase and sentence level in middle precentral gyrus (area 55b) may be preserved even in the setting of longstanding anarthria, and propose ensemble model sentence decoding using phoneme encoding in ventral precentral gyrus (area 6v) and phrase/sentence level encoding in middle precentral gyrus (area 55b) as a means to improve the accuracy of intended language decoding for participants with longstanding anarthria and locked-in syndrome.

## Methods

### Clinical Trial

Permission for this study was granted by the U.S. Food and Drug Administration and the Institutional Review Boards of Massachusetts General Hospital, Brown University, and the VA Providence Healthcare System. Research sessions were conducted with participant T17 who is enrolled in the BrainGate2 clinical trial (ClinicalTrials.gov ID: NCT00912041). All research sessions were performed at the participant’s place of residence. This manuscript does not report primary clinical trial outcomes; instead, it describes scientific and engineering discoveries that were made using data collected in the context of the ongoing clinical trial. All procedures were conducted in accordance with relevant guidelines and regulations.

### Participant T17

T17 is a man with ALS. He has tetraplegia, anarthria, and ventilator dependence; his only remaining volitional motor control is over his extraocular muscles. Following enrollment, T17 had a preoperative structural MRI, resting state fMRI, and task-based fMRI to identify appropriate anatomical targets for microelectrode array (MEA) placement. Resting state fMRI was used to generate estimated parcellations of the relevant brain areas using a custom instantiation of the Human Connectome Project ^49,50^ analysis pipeline that was modified for deployment on clinical MRI scanners able to accommodate a mechanical ventilator. Subsequently, T17 underwent placement of six 64-electrode microelectrode arrays (Blackrock Microsystems; 1.5 mm electrode length) in the left precentral gyrus; two arrays were placed in the dorsal precentral gyrus (area 6d), two arrays were placed in the ventral precentral gyrus (area 6v), and two arrays were placed in middle precentral gyrus (area 55b) (Fig. 1a). T17 consented to publication of photographs and videos containing his likeness.

### Neural Signal Processing

Voltage time series signals were recorded using the Neuroplex-E system (Blackrock Neurotech) attached to three percutaneous connectors on the participant’s head, and transmitted via three mini-HMDI cables (one to each percutaneous connector), attached to two Gemini hubs (Blackrock Neurotech), prior to final processing via a Neural Signal Processor (NSP) (Blackrock Neurotech). Neural data were streamed at 30 kHz from the NSP to our processing pipeline. Signals were analog filtered (4th order Butterworth with corners at 250 Hz to 5 kHz) using the Scipy python library (scipy.signal.filtfilt).

Linear regression referencing ^51^ (LRR) filter coefficients and subsequent electrode-specific thresholds were determined using filtered 30kHz data recorded from an initial reference block at the beginning of each session. LRR coefficients are computed by solving *Y* = *WX* where *Y* is the signal from a given electrode we require and *X* is the signal from all other electrodes. We solve for the LRR weight matrix through least squares calculation *W* : *W* = *inv*(*X^T^ X*)*X^T^ Y* where *inv* is matrix inversion ^52^. electrode specific thresholds are then calculated using filtered 30kHz data once these calculated references have been applied. Thresholds were set at −3.5 times the standard deviation of the voltage signal per electrode. The number of non-causal threshold crossing (ncTX) events were computed by counting the number of times the filtered neural time series crossed these pre-calculated thresholds. Spike band power was computed by taking the sum of squared voltages observed during each 10ms time bin. During closed-loop decoding blocks, feature normalization was employed to account for neural nonstationarities (drifts in mean firing rate) which could arise over the course of a block. Within each electrode, threshold crossing rates and spike band power were z-scored (mean subtracted and divided by standard deviation per electrode). Feature extraction (threshold crossings and spike band power), binning, decoding and task phase control were performed through the Python based, modular BRAND ^53^ framework, where each process is instantiated as a self-contained node-based python program. Messaging between these nodes is performed using a Redis database.

### Gaussian Naive Bayes

The Naive Bayes classifier is a simple probabilistic predictor which assumes conditional independence between features given the class. Gaussian Naive Bayes takes this a step further for continuous data and makes the additional assumption that values associated with each class are normally distributed. Offline single-trial classification results (reported in Fig. 1e,f,g,h,i,j and Fig. 5b,c,d,f) were generated using a cross-validated (leave-one-out) Gaussian Naive Bayes classifier. All trials are z-scored (mean subtracted and standard deviation divided) and time-averaged in the go period(s) of each task. For every trial, the classifier is trained on all other trials and evaluated on the held-out trial. We use the Scikit-learn ^54^ implementation of Gaussian Naive Bayes.

### Mahalanobis distance and hierarchical clustering

The Mahalanobis distance is a measure of the distance between a point and a distribution, which, unlike the Euclidean distance, accounts for the correlations between variables and differences in scale, making it particularly useful in multivariate statistics. Threshold crossing neural features pertaining to each stimuli are hierarchically clustered based on the Mahalanobis distance metric.

### WFST Language Model

We use a 5-gram language model implemented using a weighted finite state transducer (WFST), built on the Kaldi system ^31^. This probabilistic model was trained using the WeNet framework ^55^ and utilized the large OpenWebText2 ^32^ corpus to compute conditional probabilities. When given lattices of RNN output probability vectors, the WeNet framework initiates several Viterbi ^56,57^ searches through the most recent lattice, incorporating phoneme-by-phoneme transition probabilities when inferring the most likely typed sentence. Language model parameter details can be found in Table 2.

### Sentence Speech Decoding

Sequence decoding of attempted articulatory movements was performed using a Recurrent Neural Network (RNN) decoder trained with a Connectionist Temporal Classification (CTC) ^58–60^ loss function. This loss function allows the RNN to learn a mapping between two unaligned sequences (neural features and spoken phonemes) which are at varying temporal resolutions, similarly to recent work ^14–16^. A 5-gram language model then uses Viterbi (beam) search to infer the most likely sentence spoken given the complete sequence of RNN output probabilities at all timesteps and the statistics of the English language. Error rate, for both phonemes and words, is measured as the percent of incorrect insertions, deletions and erroneous substitutions to each phoneme when decoded in sequence.

### Recurrent Neural Network

The recurrent neural network (RNN) used for phoneme sequence decoding is implemented in Tensorflow2 ^61^ as a 5 layer gated recurrent unit (GRU) network, each with 512 units. A non-linear input layer is added per session to account for cross-session neural variability. Each non-linear layer contains the same number of units as the dimensionality of the neural features (512 features used). During training, batches of trials only from a given session are selected at random, such that the corresponding input layer is trained along with the 5 layer RNN. Training using backpropagation through time (BPTT) minimizes the Connectionist Temporal Classification loss function ^58^. Various regularization and data augmentation techniques are utilized: Dropout, Gaussian White noise, L2 weight norm. Hyperparameter details can be found in Table 1.

Much recent work focuses on reducing or eliminating the adverse decoding effects of neural nonstationarities ^62–67^. Here, we utilized training data across multiple days; a non-linear input layer is added per session to account for cross-session neural variability, as was used in recent work ^15,16,62^. Each non-linear layer contains the same number of units as the dimensionality of the neural features. During training, batches of trials only from a given session are selected at random, such that the corresponding input layer is trained along with the 5 layer RNN. This resulted in a cross-session ensemble decoder that was relatively robust to nonstationarities in the short to medium term (resulting in the reported online error rates in Fig. 3b)

### Phased Task

Participant T17 was asked to memorize a phoneme, word, phrase or sentence in the first reading phase of a phased task. In the second phase, the participant was asked to internally speak (without vocalizing) the memorized language unit. In the last phase, the participant was asked to attempt to speak with articulation the memorized language unit. A list of stimuli can be seen in Table 3. Each stimulus was repeated 5 times in a random order.

### Personalized text-to-speech model

Similarly to previous work ^16^, we use a neural network based text-to-speech model ^33^ which is first pre-trained on large spoken voice datasets such as VCTK ^68^ containing numerous utterances in English across many speakers. The model was then fine-tuned with audio recordings of the participant’s voice prior to ALS onset to produce natural and relatively faithful voicing of decoded sentences.

## Data Availability

Data relevant to this study will be publicly accessible upon publication.

## Code Availability

Code to replicate the main findings of this study will be made available in a public GitHub repository upon publication.

## Acknowledgments and Support

We thank T17, their family and carepartners for the time and effort they contributed to the BG2 trial. We thank Dr. Stephen Mernoff for his clinical monitoring of trial participants. We thank Dr. Gladys Hill for her illustrations in Figure 3. This work was supported by Office of Research and Development, Rehabilitation R&D Service, Department of Veterans Affairs (N2864C,A4820R, A2295R); NIH NIDCD (U01DC017844, K23DC021297,1DP2DC021055); NIH NINDS R25NS065743, AHA (23SCEFIA1156586); A.P. Giannini Postdoctoral Fellowship to N. Card. S.D.S has a Career Award at the Scientific Interface from the Burroughs Wellcome Fund.

Several analyses were performed using code from: https://github.com/fwillett/speechBCI.

## Author Contributions

JJJ and DBR conceived the study.

JJJ wrote the manuscript, and led the development, analysis, and interpretation of experiments.

DBR, SH and HLA reviewed analysis and interpretation of experimental data.

HH performed Mahalanobis distance and hierarchical clustering analysis.

NSC, MW, DMB, and SDS built the real-time BRAND feature extraction system, implemented the neural brain-to-text decoding pipeline in BRAND, built the task control interface, and offline analysis tools.

JJJ and AJA collected T17 speech session data.

JDS managed integration and deployment of BCI system software and hardware.

LRH is the sponsor-investigator of the multisite BrainGate2 pilot clinical trial.

ZMW, SSC, DBR and LRH planned T17’s array placement surgery, and ZMW performed T17’s array placement surgery. LRH was responsible for all clinical trial related activity at VA Providence and MGH. LRH and DBR supervised and guided all research activity with T17.

The study was supervised and guided by DBR.

All authors reviewed and edited the manuscript.

## Supplement

### Supplementary Figures

**Supplementary Figure 1:**
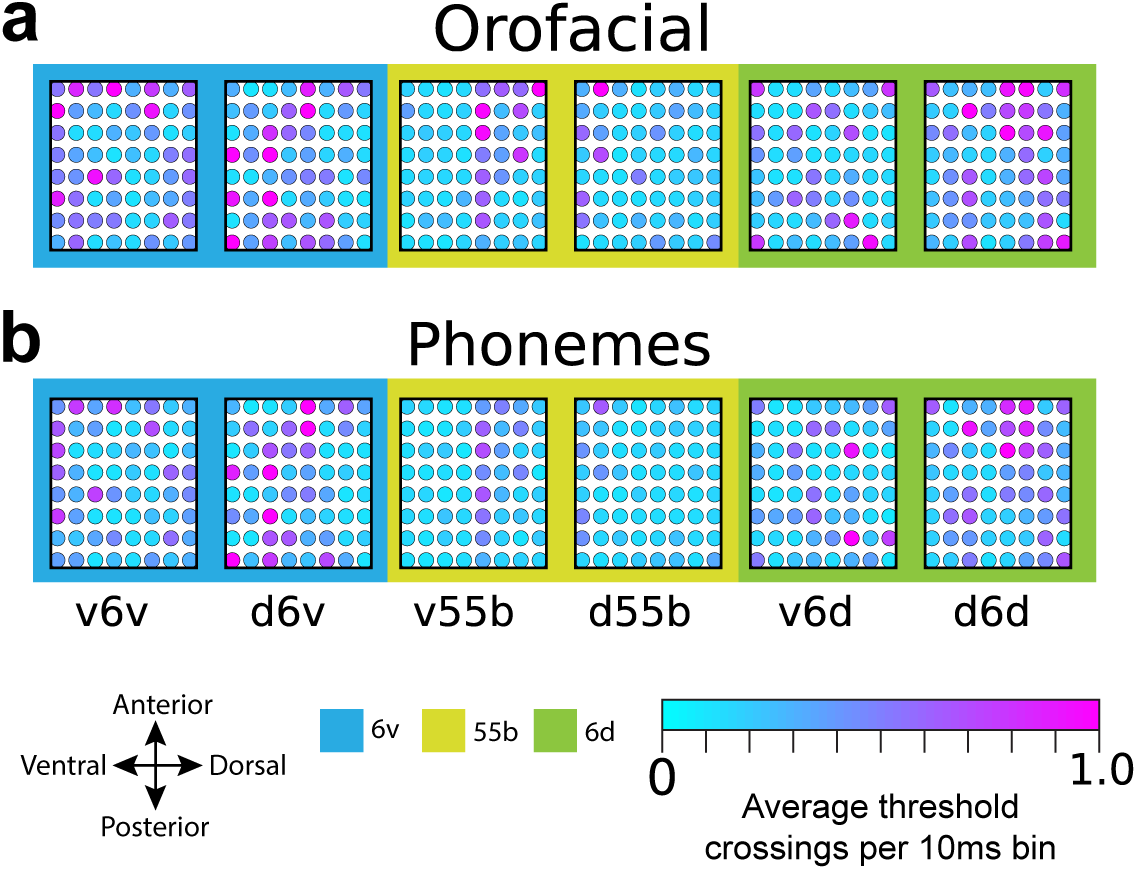
**a,b**, Average number of threshold crossings in each 10ms time bin per electrode across all 64-electrode arrays during the go periods of **a**, the isolated orofacial movements task, and **b**, the isolated phoneme task.

**Supplementary Figure 2:**
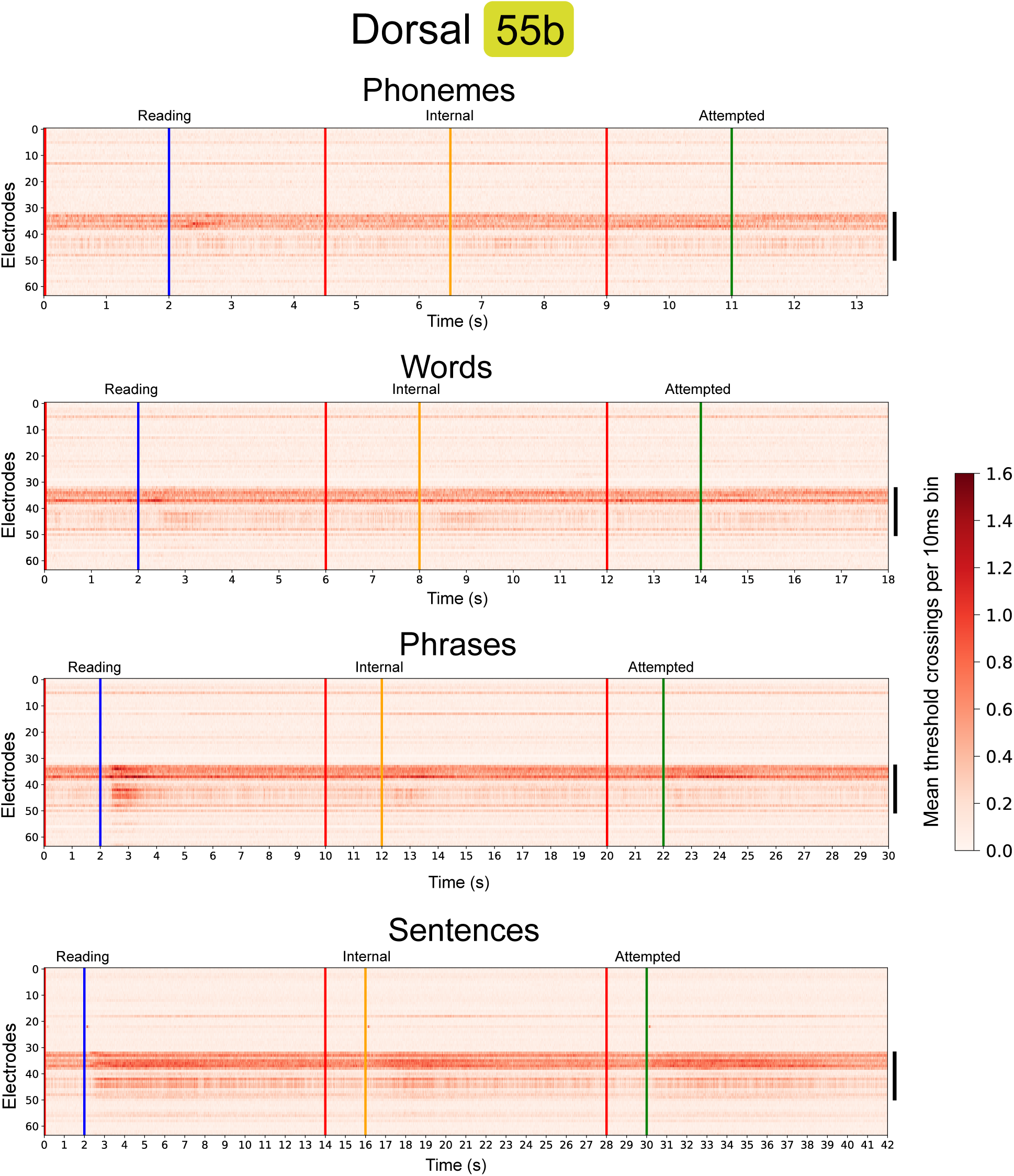
Heatmap showing average number of threshold crossings per 10ms bin across all dorsal 55b array electrodes for each language unit (phonemes, words, phrases and sentences) during all phases (reading, internal speech, attempted speech) of the phased task. Electrodes with sustained activity are marked with a black line on the right of each heatmap.

**Supplementary Figure 3:**
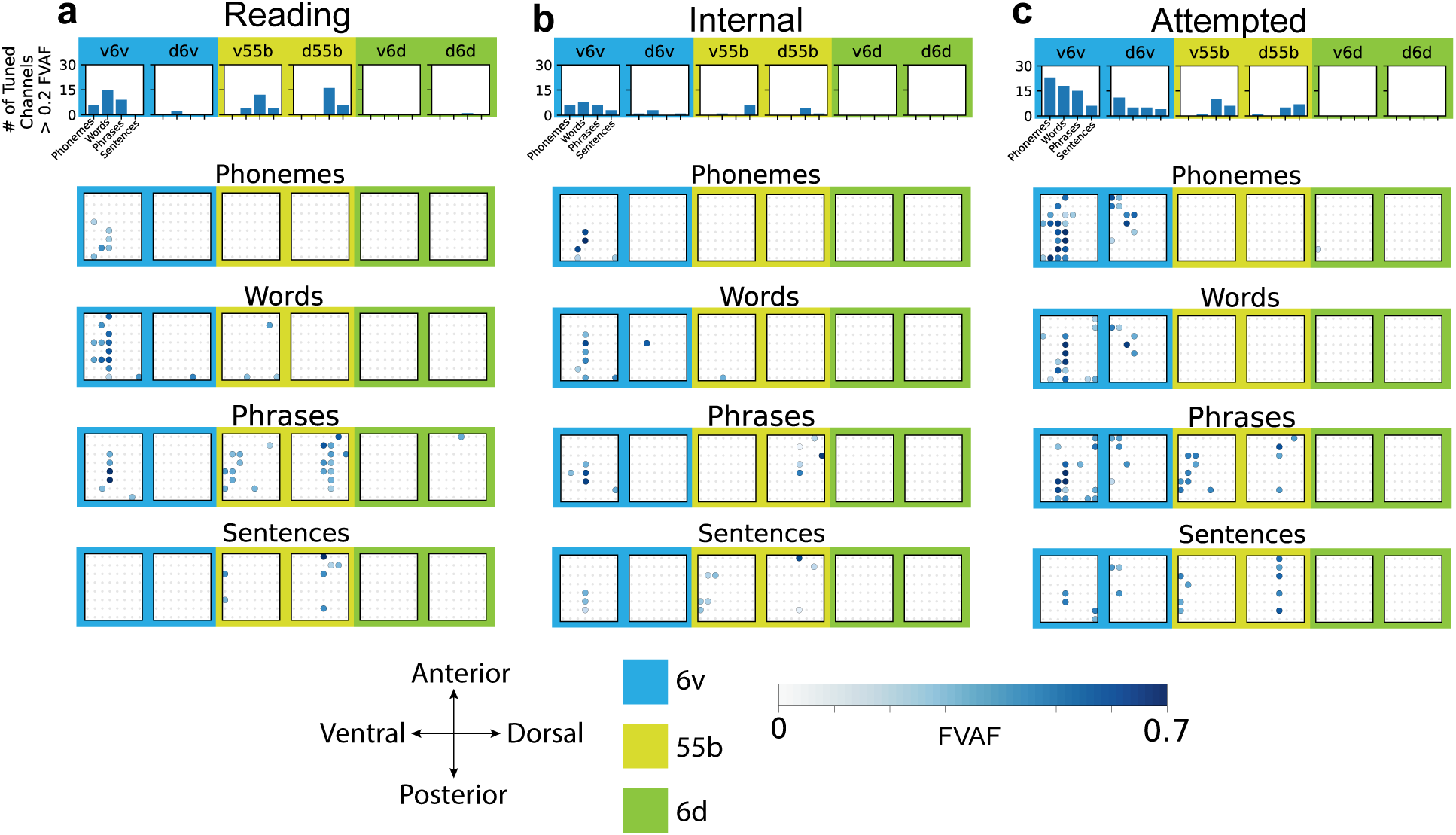
**a,b,c**, For each phase of the phased reading, internal speech and attempted speech task: Top: Number of tuned electrodes with fraction of variance accounted for above 0.2 across all electrodes of all 6 arrays for each language unit (phonemes, words, phrases and sentences). Bottom: Tuning heatmaps for all 6 arrays, with backgrounds shaded by cortical location (6v in blue, 55b in yellow, 6d in green). Circles indicate that threshold crossing features on a given electrode varied significantly across a given task phase for a given language unit (phonemes, words, phrases or sentences) (p < 1 × 10^−5^ assessed with one-way analysis of variance). Shading indicates the fraction of variance accounted for (FVAF) by within phase differences in threshold crossing rate per electrode.

**Supplementary Figure 4:**
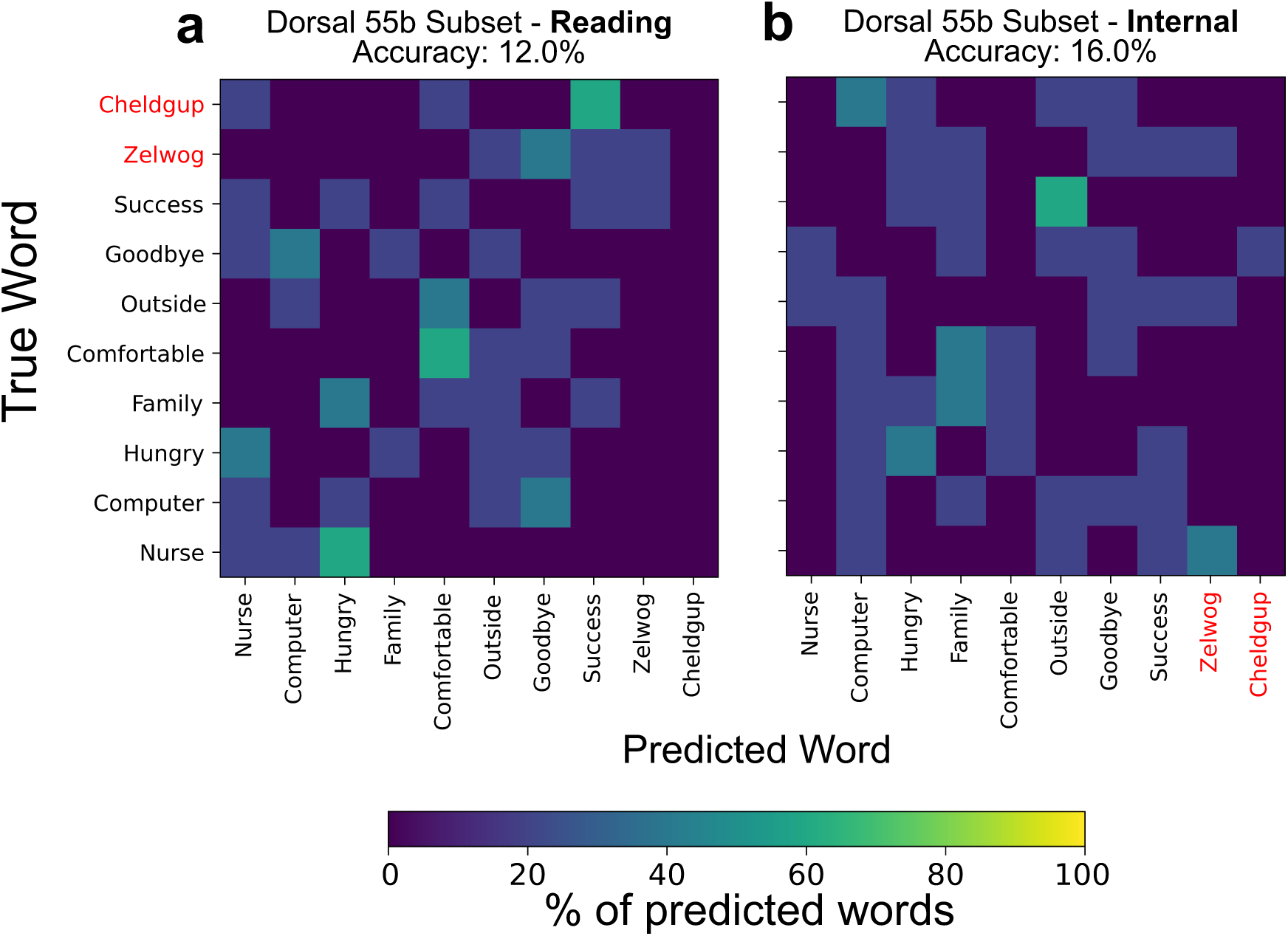
**a,b**, Confusion matrices showing decoding errors when decoding words in the **a**, reading, and **b**, imagined, phases of the task using a Gaussian Naive Bayes decoder on a subset ensemble of 20 contiguous electrodes from the dorsal 55b array. Nonsense word stimuli are highlighted in red.

### Supplementary Tables

**Supplemental Table 1:** RNN decoding model parameters.

| Parameter Name | Parameter Description | Value |
| --- | --- | --- |
| # Units | Units in each GRU layer | 512 |
| # Layers | Number of GRU layers | 5 |
| Kernel Size | Number of feature time bins input to the RNN model (size of sliding window) | 14 |
| Kernel Stride | Size of sliding window shift | 4 |
| L2 Regularization | Backpropagation penalizes large RNN weight sizes | 0.00001 |
| Dropout | Probability of RNN units being excluded from the network during training | 0.4 |
| Input layer # Units | Number of units in each session specific input layer | 512 |
| Input layer Dropout | Probability of input layer units being excluded from the network during training | 0.2 |
| Input activation | Input layer activation function | softsign |
| White noise SD | Standard deviation of Gaussian (zero mean) noise added to neural features (data augmentation) | 1.5 |
| Constant offset SD | Standard deviation of Gaussian (zero mean) noise added to all timesteps of neural features (data augmentation) | 0.2 |
| Smooth Kernel SD | Standard deviation of temporal Gaussian smoothing kernel applied to neural features | 2 |
| Batch Size | Number of sentences used to train the RNN model per step (all taken from same session in a single batch) | 64 |
| Training batches | Number of training batches (training ended early if error rate did not improve for 500 iterations) | 2,000 to 10,000 |
| Learning rate | Linearly decaying learning rate | 0.02 to 0 |
| $\beta_1$ | The exponential decay rate for the 1st moment estimates for ADAM optimizer | 0.9 |
| $\beta_2$ | The exponential decay rate for the 2nd moment estimates for ADAM optimizer | 0.999 |
| $\epsilon$ | Constant for numerical stability for ADAM optimizer | 0.1 |

**Supplemental Table 2:** 5-gram Language model parameters.

| Parameter Name | Parameter Description | Value |
| --- | --- | --- |
| Max Active | Maximum active states during beam search | 7000 |
| Min Active | Minimum active states during beam search | 200 |
| Beam | Beam length | 17 |
| Lattice Beam | Desired search through RNN logit lattice | 8 |
| Acoustic Scale | Weighting of Language model compared to RNN logit probabilities, lower value weights RNN more | 0.4 |
| CTC blank skip threshold | RNN logits with a blank probability exceeding this threshold are skipped during inference | 1.0 |
| Length Penalty | Negative weighting of sequence length | 0.0 |
| nBest | Number of sentences to infer in order of likelihood | 100 |
| Blank Penalty | Negative weighting applied to blank class | 4 |
| Rescore | Boolean determining G.fst rescoring of nBest inferred sentences | False |

**Supplemental Table 3:** Phase Task Cues.

| Phonemes | Words | Phrases | Sentences |
| --- | --- | --- | --- |
| Pah | Nurse | I like my music. | All animals are equal, but some are more equal than others. |
| Rah | Computer | They have tired faith. | Spring was moving in the air above and in the earth below. |
| Dah | Hungry | You need clean glasses | The human eye can distinguish over ten million different colors. |
| Ah | Family | What is right here? | The conch shell exploded into a thousand white fragments. |
| Shah | Comfortable | My family is outside. | Hydrogen is the most abundant element in the known universe. |
| Lah | Outside | Bring it up closer. | Magna Carta sought to prevent the king from exploiting power. |
| Tah | Goodbye | Tell my nurse goodbye. | Knock-kneed, coughing like hags, we cursed through sludge. |
| Oy | Success | That is not how it is. | Every man desires to live long, but no man wishes to be old. |
| Hah | Zelwog | Faith hungry computer. | Cytoplasm is the gelatinous liquid that fills the inside of a cell. |
| Gah | Cheldgup | Need how yes feel. | Reaching the South Pole in the heroic age of Antarctic exploration. |

**Supplemental Table 4:** T17 Session Data.

| **Trial Day** | **Tasks** | **Number of Sentences** |
| --- | --- | --- |
| 26 | 50-word Vocabulary Sentences | 200 |
| 33 | 50-word Vocabulary Sentences<br>125,000-word Vocabulary Sentences | 480 |
| 46 | 125,000-word Vocabulary Sentences | 360 |
| 47 | 125,000-word Vocabulary Sentences | 200 |
| 48 | 125,000-word Vocabulary Sentences | 240 |
| 67 | 125,000-word Vocabulary Sentences | 280 |
| 68 | 125,000-word Vocabulary Sentences | 320 |
| 74 | 125,000-word Vocabulary Sentences | 80 |
| 75 | 125,000-word Vocabulary Sentences | 240 |
| 89 | 125,000-word Vocabulary Sentences | 160 |
| 94 | 125,000-word Vocabulary Sentences | 40 |
| 95 | 125,000-word Vocabulary Sentences | 320 |
| 164 | 50-word Vocabulary Sentences | 360 |
| 165 | 50-word Vocabulary Sentences | 360 |
|  |  | **Total**<br>3640 |

